# Movement and conformity interact to establish local behavioural traditions in animal populations

**DOI:** 10.1101/338657

**Authors:** Marius Somveille, Josh A. Firth, Lucy M. Aplin, Damien R. Farine, Ben C. Sheldon, Robin N. Thompson

**Affiliations:** Edward Grey Institute, Department of Zoology, University of Oxford, South Parks Road, Oxford OX1 3PS, UK.; Max Planck–Yale Center for Biodiversity Movement and Global Change, Yale University, 165 Prospect Street, New Haven, CT 06520-8105, USA.; Merton College, Merton Street, University of Oxford, Oxford, OX1 4JD; Cognitive and Cultural Ecology Research Group, Max Planck Institute for Ornithology, Radolfzell, Germany.; Department of Collective Behaviour, Max Planck Institute for Ornithology, Universitätsstrasse 10, 78457 Konstanz, Germany.; Chair of Biodiversity and Collective Behaviour, Department of Biology, University of Konstanz, Universitätsstrasse 10, 78457 Konstanz, Germany.; Department of Zoology, University of Oxford, South Parks Road, Oxford OX1 3PS, UK.; Mathematical Institute, University of Oxford, Radcliffe Observatory Quarter, Woodstock Road, Oxford OX2 6GG, UK.; Christ Church, University of Oxford, St. Aldates, Oxford OX1 1DP, UK.

## Abstract

The social transmission of information is critical to the emergence of animal culture. Two processes are predicted to play key roles in how socially-transmitted information spreads in animal populations: the movement of individuals across the landscape and conformist social learning. We develop a model that, for the first time, explicitly integrates these processes to investigate their impacts on the spread of behavioural preferences. Our results reveal a strong interplay between movement and conformity for determining whether local traditions establish across a landscape or whether a single preference dominates the whole population. The model is able to replicate a real-world cultural diffusion experiment in great tits *Parus major*, but also allows for a range of predictions for the emergence of animal culture under various initial conditions, habitat structure and strength of conformist bias to be made. Integrating social behaviour with ecological variation will be important for understanding the stability and diversity of culture in animals.

## INTRODUCTION

The social transmission of information plays a central role in the lives of many animal species [1–3]. Social learning via observation of, or interaction with, other individuals is an efficient mechanism for acquiring information about the environment, leading to adaptive adjustments of behavioural responses [4,5]. The transmission of information through social networks can lead to the emergence of regional variations in behaviour that are stable through time (called local cultures or traditions; [6–10]). However, we still have little mechanistic understanding of the conditions under which local cultures can emerge. Understanding how ecological, cognitive and social processes determine the spread of information between individuals in wild populations is crucial if we want to discern the conditions under which information spreads and local traditions emerge.

A key ecological process that is likely to affect the spread of information is movement. First, movement of animals between discrete groups or sub-populations is expected to accelerate information spread across the whole population. Second, moving individuals can potentially import different behaviours into local groups or sub-population [11,12]. How individuals move in a landscape, itself influenced by a range of factors such as habitat structure [13] and demography [14], is therefore likely to shape the dynamics of behaviours in natural populations.

One of the main socio-cognitive factors thought to affect the emergence of culture is conformity [15]. Conformist social learning is here defined as positive frequency copying, where individuals are disproportionately likely to adopt the most common behavioural trait [16]. Importantly, if individuals exhibit conformist learning, a single socially learnt behavioural preference might fix in a group, remain stable over time and be resistant to invasion by alternative variants, leading to the establishment and persistence of group-specific traditions. Several experimental studies have therefore suggested that conformity plays an important role in the establishment and stability of local traditions in various animal species [11,12,17,18]. However, the interplay between social learning biases such as conformity, and the ecological factors that determine the context in which such learning takes place, has rarely been studied.

Here, we investigate how movement and conformity interact to shape the spread of socially-transmitted information and the establishment of local traditions in animal populations. We first develop a theoretical spatially-explicit model of the spread of behavioural preference in which a puzzle (representing a novel foraging resource) can be solved in one of two ways. The population is assumed to be composed of several spatially distinct sub-populations, and each individual can either be unable to solve the puzzle, or solve the puzzle and prefer one of the two solutions. Individuals can learn the behaviour from each other, with a conformist bias guiding which preference they learn, and move between sub-populations. We then use this model to investigate the conditions under which local cultural traditions emerge. We consider simple scenarios in which there are only two or three sub-populations. In addition, we test the model’s ability to replicate a real-world cultural diffusion experiment [11], in which alternative novel foraging techniques – consisting of opening a bi-directional door puzzle-box by sliding it either left or right to access food – were introduced in several wild sub-populations of great tits *Parus major*. The spread of these foraging behaviours in the population was monitored, revealing that the behaviour was socially transmitted with a significant conformist bias, and that local foraging traditions established within the population. As we show, our modelling approach can recreate the empirical results of the experiment conducted by Aplin et al. [11], and predict the conditions under which such local traditions are likely to establish and persist in the population.

## RESULTS

### The baseline model

Our model integrates the movement of individuals across the landscape and the social process of transmission of information between individuals, including a conformist bias. In the baseline version of the model, the spread of behavioural preferences for solutions to a puzzle (solutions s_1_ and s_2_) occurs in an environment with a metapopulation structure composed of two connected patches, each one hosting a sub-population (see details in Materials and Methods). These sub-populations are composed of naïve individuals (i.e. individuals who do not know how to perform the behaviour whose spread is being modelled) and knowledgeable individuals, or innovators (i.e. individuals who know how to perform one of the behaviours). At the start of each numerical solution of the model (hereafter, “simulation”), innovators are introduced in the system (initially only comprised of naïve individuals), with different behavioural preferences in each sub-population. The spread of behavioural preferences is then simulated for 150 time-steps.

We found that the emergence of contrasting local traditions (i.e. the situation where, at the end of the simulation, one sub-population is dominated by individuals with one behavioural preference while the other sub-population is dominated by individuals with the alternative behavioural preference) strongly depends on the relationship between the strength of conformity and the rate of movement of individuals between patches, determined by parameters *λ* and *m* respectively (see details in Materials and Methods). When conformity was not included in the model (i.e. *λ* = 1), the pattern that always emerged regardless of other parameter values (except when the difference in size between the two sub-populations was very large) was a mixture of behavioural preferences in both patches (Figs 1 and S1). When conformity was strong relative to the movement rate, local traditions established and remained stable (Fig 1). When conformity was weak relative to the movement rate, one of the two behavioural preferences dominated the whole system (Fig 1). In the latter case, which preference dominated depended on the relative sizes of the sub-populations. When the size of the sub-population (i.e. the number of naïve individuals in the patch) in which an innovator with a given behavioural preference was initially introduced was larger than the size of the other sub-population, then that preference came to dominate at the end of the simulation (Fig S1). When the two sub-populations had exactly the same size, which behavioural preference dominated at the end of the simulation appeared to be random and very sensitive to the precise parameter values (Fig S1). Finally, the main effect of the learning rate was to determine how much stronger/weaker conformity must be relative to the magnitude of the movement rate to obtain these patterns: the lower the learning rate, the stronger conformity had to be relative to the movement rate to see local traditions establish and stabilise (Figs 1 and S1).

**Fig 1.**
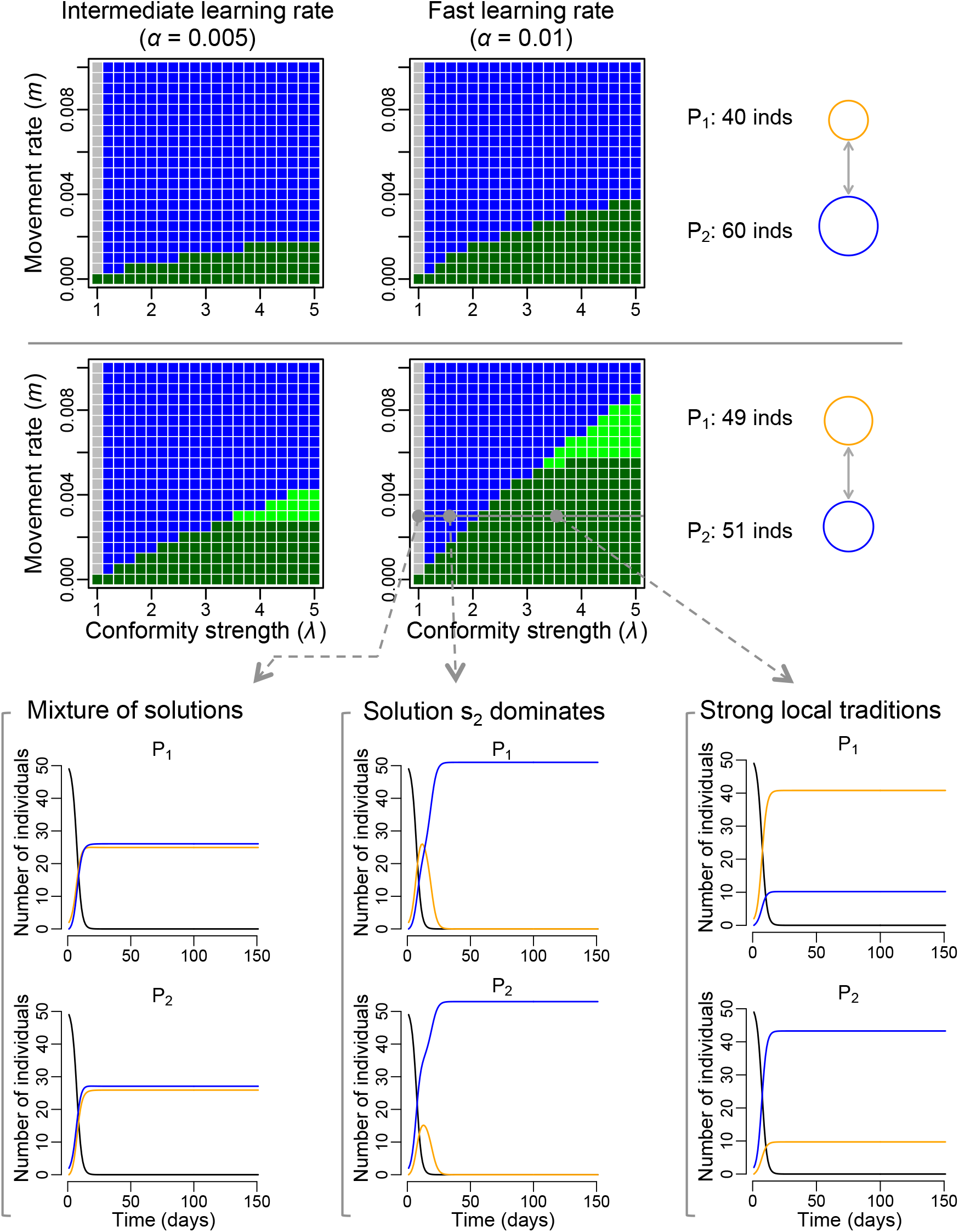
Local traditions emerge when conformity is strong relative to the movement rate. This panel shows the model outputs for the baseline model with two patches. At the start of every simulation, patch 1 contained two innovators using solution s_1_ and contained P_1_ naïve individuals, while patch 2 contained two innovators using solution s_2_ and contained P_2_ naïve individuals. Each pixel in the phase diagrams corresponds to a simulation run with the corresponding parameter values, and the colour of the pixel indicates the emerging pattern after 150 days: grey = mixture of solutions in every patch, blue = solution s_2_ dominated the whole system, light green = weak local traditions, dark green = strong local traditions. The two columns of phase diagrams represent different learning rates: intermediate (*α* = 0.005) and fast (*α* = 0.01). The two rows of phase diagrams represent a different configuration of patch sizes, reflecting the differences in number of naïve individuals at the start of the simulation (U_0_) occurring in each patch. Three examples of the evolution of the number of naïve individuals (black curve) and number of solvers using solution s_1_ (orange curve) and solution s_2_ (blue curve) in each patch, are shown for fixed movement rate (m = 0.003), learning rate (*α* = 0.01) and patch sizes configuration, but with varying conformity strength. When no conformity bias was included (*λ* = 1), patches contained a mixture of solutions; when conformity was relatively weak (*λ* = 1.5), solution s_2_ (that seeded in the larger population) ended up dominating in both patches; and when conformity as relatively strong (*λ* = 3.5), local traditions emerged.

### Simple environmental setting

To examine the role of space in the spread of behavioural preferences, we extended the baseline model to a simple environmental setting with three spatially distinct patches, thereby effectively adding an extra patch containing no innovators at the start of the simulation (see Materials and Methods for details). Once again, when conformity was not included in the model (*λ* = 1), the pattern that always emerged regardless of other parameter values was a mixture of behavioural preferences in all patches (Fig 2). Similarly to the baseline model, there was a strong relationship between movement rate and conformity strength; when conformity was strong relative to the movement rate, local traditions established and were stable (i.e. two sub-populations were dominated by individuals with one behavioural preference while the third sub-population was dominated by individuals with the alternative behavioural preference; Fig 2), and when conformity was weak relative to the movement rate, one of the two behavioural preferences ultimately dominated the entire system (Fig 2). In the latter case, which preference dominated depended on both the relative sizes of the sub-populations as well as the spatial configuration (i.e. the distances separating the sub-populations, which determines the relative movement rates of individuals between pairs of sub-populations). If the size of the sub-population in which innovators with a given behavioural preference were initially introduced was larger than the size of the sub-population in which innovators with the alternative behavioural preference were introduced, and all sub-populations were equidistant, then the former preference was predominant at the end of the simulation (Fig S2). Also, increasing the distance separating a sub-population in which innovators with a given behavioural preference were initially introduced from the two other sub-populations (i.e. creating unequal movement rates between patches) resulted in this preference not being able to dominate the system at the end of the simulation (Fig S2).

**Fig 2.**
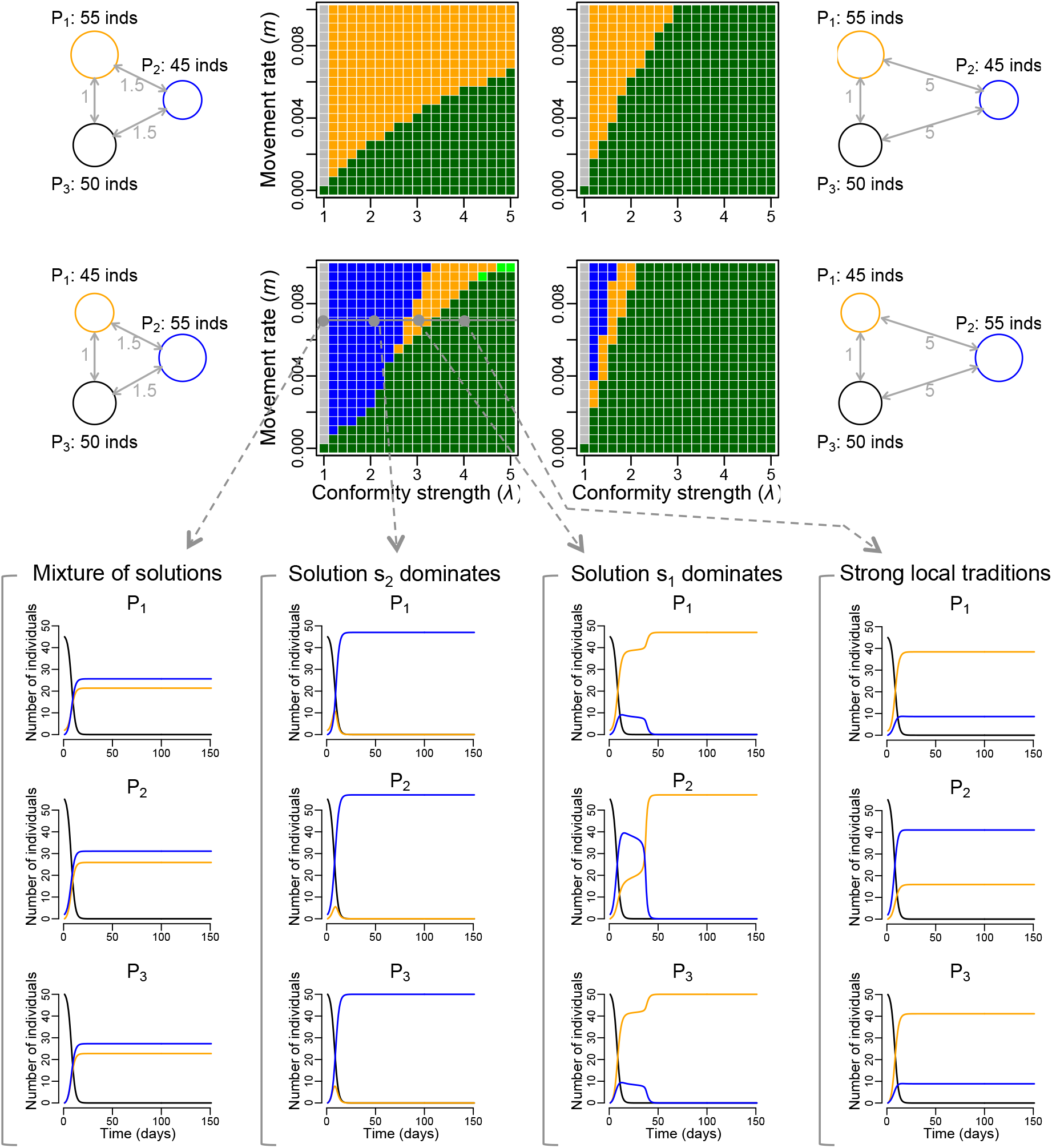
Space has a complex effect on how conformity and movement lead to either the emergence of local traditions or the domination of a single solution. This panel shows the model outputs in the three-patch case. At the start of every simulation, patch 1 contained two innovators using solution s_1_ and contained P_1_ naïve individuals, while patch 2 contained two innovators using solution s_2_ and contained P_2_ naïve individuals, and patch 3 contained only naïve individuals. Each pixel in the phase diagrams corresponds to a simulation run with the corresponding parameter values, and the colour of the pixel indicates the emerging pattern after 150 days: grey = mixture of solutions in every patch, blue = solution s_2_ dominated the whole system, orange = solution s_1_ dominated the whole system, light green = weak local traditions, dark green = strong local traditions. The two columns of phase diagrams represent different spatial configurations: a different distance between patch 2 and the other two patches (whose distance separating them was set to 1), which results in different relative migration rates between pairs of patches. The two rows of phase diagrams represent a different configuration of patch sizes, reflecting differences in the number of naïve individuals (inds) at the start of the simulation (U_0_) that occurred in patches P_1_ and P_2_. Four examples of the evolution of the number of naïve individuals (black curve) and number of solvers using solution s_1_ (orange curve) and solution s_2_ (blue curve) in each patch, are shown for fixed movement rate (m = 0.007), and spatial and patch sizes configuration, but with varying conformity strength. When no conformity bias was included (*λ* = 1), patches contained a mixture of solutions; when conformity was relatively weak (*λ* = 2), solution s_2_ ended up dominating in every patch; when conformity was intermediate (*λ* = 3), solution s_1_ ended up dominating in every patch; and when conformity was relatively strong (*λ* = 4), local traditions emerged.

Increasing the distance between patches also affected how much stronger/weaker conformity must be in comparison to the movement rate to generate the emerging patterns: the smaller the distance, the stronger conformity had to be relative to the movement rate for local traditions to establish and stabilise (Figs 2 and S2). When both patch size and spatial configuration acted in opposite directions, then either behavioural preference could ultimately dominate depending on the relationship between the strength of conformity and the magnitude of the movement rate. This may arise when the size of the sub-population in which innovators with a given behavioural preference were initially introduced was larger than the size of the sub-population in which innovators with the alternative behavioural preference were initially introduced, but this sub-population was also further away form the two others (see illustration in Fig 2). When conformity was relatively strong, but not so strong as to generate local traditions, then the behavioural preference that was not released in the larger, further away sub-population ended up dominating the system, otherwise the alternative preference dominated at the end of the simulation (Fig 2).

Unexpected results were also observed when both sub-populations in which innovators were introduced at the start of the simulation had the same size, which is larger than the third sub-population, with one of these two sub-populations being slightly further away from the two others. In contrast to what might have been expected based on other results, when conformity was very weak compared to the movement rate, then the behavioural preference of the innovator that was initially introduced in the most distant sub-population ended up being the most prevalent in the system (Fig S2, second plot of fourth row, in blue).

### Realistic environmental setting

To examine the role of habitat structure and the ecological process of movement in a realistic setting, we extended the baseline model to represent the great tit population of Wytham Woods, which has been the subject of a long-running study, and the site of a recent cultural diffusion experiment [11]. Running the model of spread of behavioural preference for this real-world animal population in its natural environment (see Materials and Methods for details) yielded results that were consistent with those for the baseline model and its extension to three patches. Three possible patterns emerged at the end of the simulation depending on parameter values: (1) a mixture of behavioural preferences in every sub-population when conformity was not included in the model (i.e. *λ* = 1), (2) domination of one behavioural preference across the population when conformity was weak relative to the magnitude of the movement rate, and (3) the establishment of local traditions when conformity was strong relative to the magnitude of the movement rate (i.e. some sub-populations were dominated by individuals with one behavioural preference while the rest were dominated by individuals with the alternative preference, Fig 3). Increasing the degree of fragmentation of the landscape (by only allowing individuals to move through contiguous forest, as opposed to moving along straight direct paths between patches) affected how much stronger/weaker conformity must be compared to the magnitude of the movement rate to generate the different emerging patterns (Fig 3). Increasing the learning rate had a similar effect to increasing the degree of landscape fragmentation (Fig 3). That is, for local traditions to establish and stabilise, conformity had to be stronger relative to movement rates when either the habitat was less fragmented or when the learning rate was slower (Fig 3).

**Fig 3.**
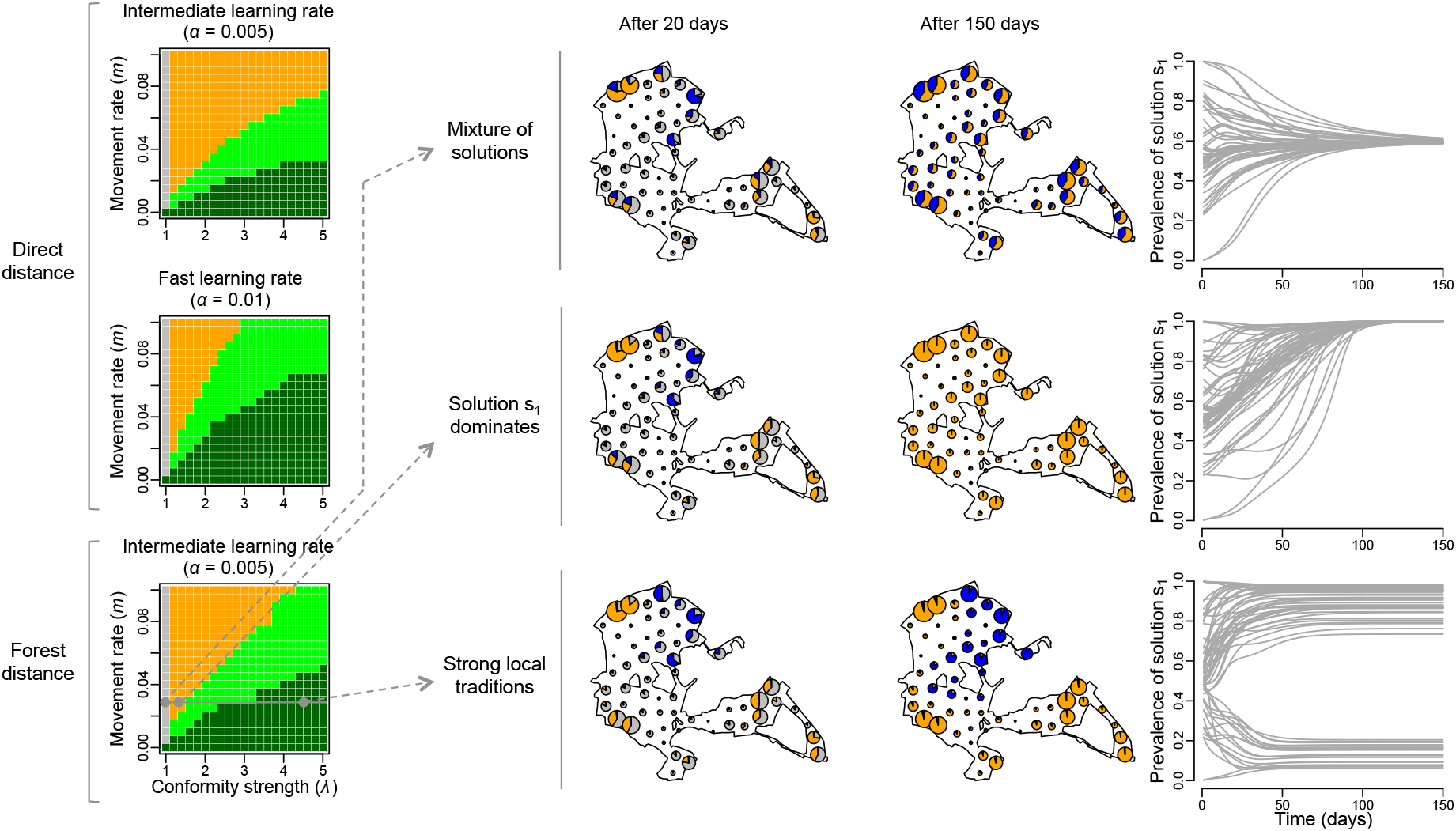
Predictions for the spread of information in great tit cultural diffusion experiment in Wytham Woods. This analysis was designed to replicate the conditions of the cultural diffusion experiment performed by Aplin et al. [11], in which two alternative foraging techniques were introduced in the great tits population of Wytham Woods in the United Kingdom, and their spread through time monitored. Each pixel in the phase diagrams corresponds to a simulation run with the corresponding parameter values, and the colour of the pixel indicates the emerging pattern after 150 days: grey = mixture of solutions in every patch, blue = solution s_2_ dominates the whole system, orange = solution s_1_ dominates the whole system, light green = weak local traditions, light green = strong local traditions. The two top phase diagrams (with intermediate and fast learning rates) correspond to model runs in which the distance separating patches is the Euclidean distance (direct distance), while the phase diagram at the bottom corresponds to model runs using the shortest route through the forest separating pairs of patches (forest distance). Three examples of the evolution of the prevalence of solution s_1_ among solvers are shown for fixed movement rate (m = 0.03), learning rate (*α* = 0.005) but with varying conformity strength. The maps represent the extent of Wytham Woods and the location of the 60 feeders. Each feeder is represented by a pie chart indicating the number of naïve individuals (grey), solvers using solution s_1_ (orange) and solvers using solution s_2_ (blue), and the size of the pie chart is proportional to the total number individuals occurring around the feeder. When no conformity bias is included (*λ* = 1) it results in a mixture of solutions in every patch; when conformity is relatively weak (*λ* = 1.3) solution s_1_ ends up dominating in every patch; and when conformity is relatively strong (*λ* = 4.5) local traditions emerged.

The emergence of patterns depended on precisely where innovators with preferences for solutions s_1_ and s_2_ occurred, particularly when conformity was weak relative to the magnitude of the movement rate (Fig 4). When this was the case (i.e. *λ* = 1.2 and *m* = 0.02), 62% of simulations in which initial conditions were randomised resulted in the emergence of local traditions, and the rest of the simulations resulted in one behavioural preference dominating the whole system (with half of those simulations leading to solution s_1_ to be predominant, and the other half of simulation with solution s_2_ dominating). Which behavioural preference ended up dominating was strongly affected by the sizes of the pools of naïve individuals in contact with innovators preferring each solution at the start of the simulation. If one behavioural preference came to dominate the whole system, then it was likely to be the preference that was initially added in the comparatively larger sub-population (Fig 4), consistent with previous results. However, when conformity was strong relative to the magnitude of movement rate (i.e. *λ* = 4 and *m* = 0.02), all the simulations resulted in the emergence of local traditions. When no conformity was included (i.e. *λ* = 1 and *m* = 0.02), 79% of simulations resulted in a mixture of behavioural preferences in every sub-population, and the rest of simulations resulted in one behavioural preference dominating the whole system (with half of simulations leading to solution s_1_ throughout the landscape, and the other half of simulations with solution s_2_ predominant). Once again, the sizes of the pools of naïve individuals in contact with each innovator at the start of the simulation affected which pattern emerged, similarly to when a weak conformity was included (Fig 4).

**Fig 4.**
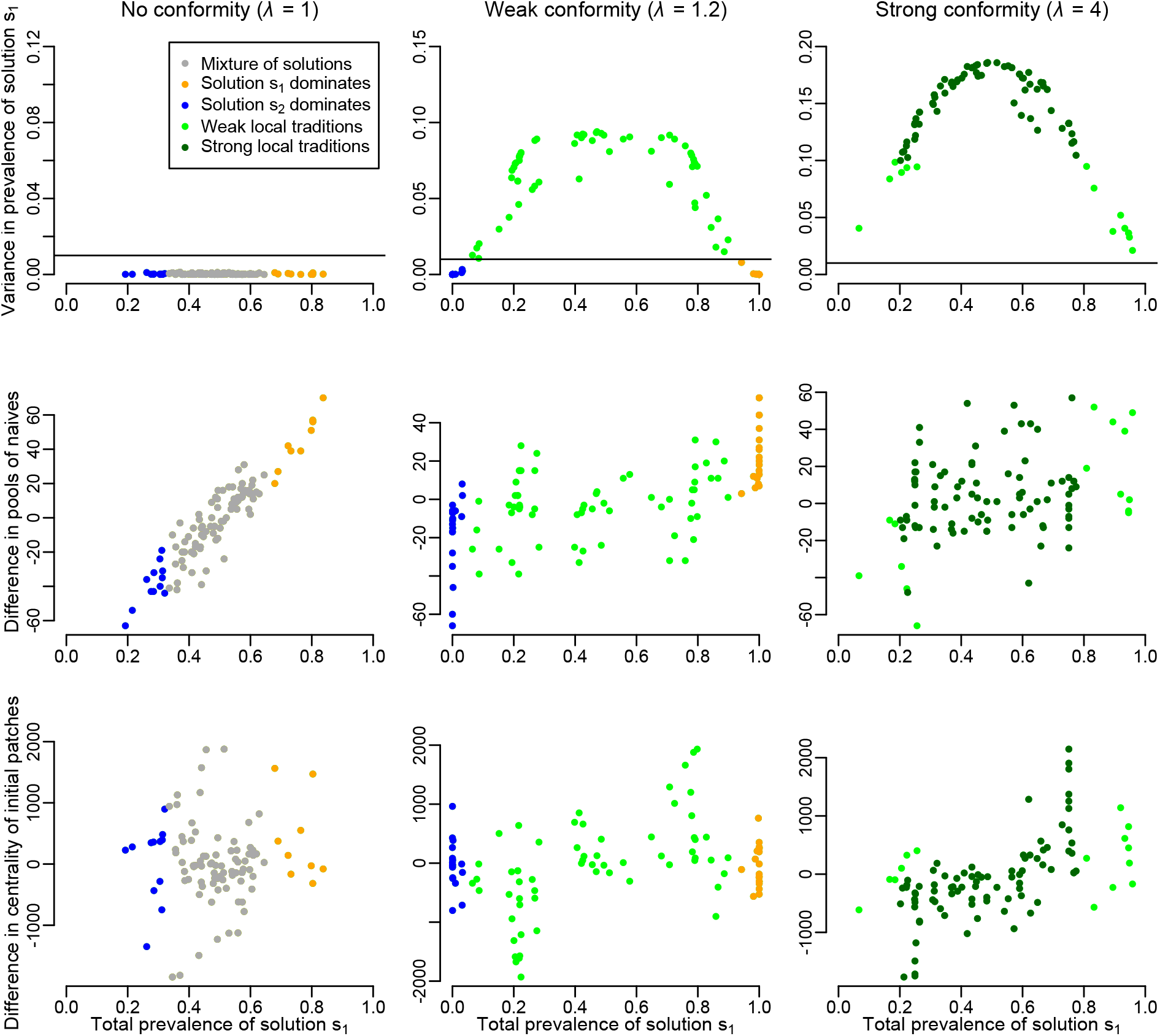
The outcome of the spread of information is sensitive to the initial conditions. This panel shows results for the randomisation of initial conditions for the spread of information in Wytham Woods. Each column of plots corresponds to a different conformity strength, for which 100 simulations with randomised initial conditions were run. The first row of plots indicates the values of simulations for the two summary statistics used to identify the emerging pattern. The horizontal line indicates the threshold above which local traditions were said to have emerged. The second row of plots investigates a relationship between the total prevalence of solution s_1_ (one of the two summary statistics) and the difference between the pool of naïve individuals initially in contact with solution s_1_ and s_2_ (i.e. size of the sub-population in which innovators with solution s_1_ were released at the start of the simulation minus the size of the sub-population in which innovators with solution s_2_ were released). The third row of plots investigates a relationship between the total prevalence of solution s_1_ and the difference in the centrality of the sub-populations in which innovators with solutions s_1_ and s_2_ were released at the start of the simulation. The centrality of a sub-population was computed as the median distance between itself and other sub-populations (the smaller the value the more central is the sub-population). The smaller the difference in centrality, the more solution s_1_ was released in a central location compared to solution s_2_.

## DISCUSSION

Our results demonstrate the importance of the relationship between movement and conformity for determining whether or not local traditions establish in animal populations. First, our model indicates that a conformist bias in learning is key for the emergence of local traditions, as none of our simulations in which a conformist bias was not included led to the generation of local traditions (Figs 1–4; except in the two-patch case when the difference in size between the two sub-populations is very large and the learning rate is relatively fast, Fig S1). The importance of conformity in this scenario is in line with previous hypotheses and indications from experimental results [11,12,16,18]. Second, we extended this finding to show that local traditions establish only when conformity is relatively strong compared to the magnitude of the movement rate of individuals between sub-populations. This was observed for the baseline model (two patches; Fig 1) and its extension to three patches (Fig 2) as well as for the realistic environmental setting of Wytham Woods (Fig 3). As highlighted for the baseline model, when conformity was weak relative to the magnitude of the movement rate, moving individuals could continuously invade a patch with alternative behavioural preferences at a faster rate than which they could conform to the local behavioural preference in that patch, thereby leading to the domination of a single behavioural preference across the whole system by the end of the simulation (e.g. Figs 1–3). Since neither of the two alternative behavioural preferences had a selective advantage, the solution that ended up dominating was determined by the initial conditions: a given behavioural preference that started in a larger pool of naïve individuals was more likely to dominate (Figs 1, 2 and 4). This is because it spread more quickly at the start of the simulation than the alternative preference, and knowledgeable individuals moving out of that sub-population therefore represented a relatively large proportion of knowledgeable individuals in each sub-population that they arrived in.

Importantly, the spatial configuration of patches also influences the outcome of the spread of information. Increasing habitat fragmentation led to more favourable conditions for the establishment of local traditions by lowering the movement rate and thus increasing the relative impact of conformist learning. This was observed for both the three-patches setting (Fig 2) and in the realistic setting of Wytham Woods (Fig 3; when we used *forest distance*, the fragmentation of the habitat was effectively larger than when we used *Euclidean distance*). The spatial configuration of patches also affects which of the two alternative behavioural preferences ultimately dominates when conformity is weak relative to the magnitude of the movement rate. In the three-patch setting, the behavioural preference that colonised the third patch first (in which no innovators were introduced) was generally the one that ended up dominating at the end. This effect was determined by how far the patches were from each other and the relative sizes of the sub-populations. Interestingly, if the sub-population with the largest pool of naïve individuals at the start of the simulation was also more distant from the other sub-populations, which preference predominated at the end depended on the interplay between conformity strength and the magnitude of movement rate (Fig 2 second row of phase diagrams), as this affected which preference was better at colonising the patch with no innovator.

Finally, a surprising effect of space was observed in a three-patch landscape in which innovators were initially introduced in two large patches but where one of the two large patches was located slightly further away from the other two patches. In this case, the behavioural preference of the innovator in the most distant patch ended up dominating the whole system when conformity was very weak relative to the magnitude of the movement rate (Fig S2, second plot of fourth row, in blue). A possible explanation for this result is that, with a very high movement rate relative to conformity strength, naïve individuals from the patch without innovators moved *en masse* and slowed down the initial spread of the behavioural preferences. This effect was less pronounced for the preference introduced in the most distant patch (as movement was dependent on distance) and so this preference could subsequently colonise the patch without innovators faster than the alternative preference. Overall, these results highlight the important effects of habitat configuration and fragmentation on the spread of culture in animal populations (see also [19]), and allow for testable predictions to be made. It is particularly relevant given the wide range of animal populations around the world that are affected by habitat fragmentation [20,21].

Our model replicated the diffusion curves empirically observed in a cultural diffusion experiment in great tits in Wytham Woods [11]. With the same initial conditions in our model as in the field experiment (i.e. trained innovators released at the same locations in the landscape) and for an intermediate learning rate, the model predicted that sub-populations in which trained individuals were released should have reached a proportion of solvers approximately between 0.6–0.8 after 20 days (Fig S3; with one exception with a proportion of solvers of 0.35), and sub-populations in which no trained innovators were released to contain a proportion of solvers between 0.1–0.4 after 20 days (Fig S3). These values are very similar to the empirical results reported in the original study (Fig 1b in [11]). This supports the potential for this model to be used to make predictions about when novel behaviours could result in local traditions. These predictions could in turn be tested in cultural diffusion experiments.

Our model predicts that local traditions establish when the movement rate of individuals between sub-populations is low relative to the strength of conformity, based on a given learning rate. If movement rates are relatively high, the location where the different behaviours emerge has an important effect on the outcome. The model predicts that the centrality of location in the landscape largely does not affect the outcome but that the size of the pool of naïve individuals living there has a strong effect (Fig 4). If a conformist bias exists in the transmission of information, and for a given movement ability of the population, local traditions are more likely to establish and be well pronounced if two different behavioural preferences appear in sub-populations with similar sizes (Fig 4). These predictions have many implications for studying the emergence of behavioural traditions in animal populations in which social learning occurs. They highlight the key and often neglected role of movement, and particularly its interplay with conformist learning, as well as the importance of the initial conditions. It should therefore be interesting going forwards to test these model predictions for species with different levels of mobility – e.g. high mobility of fission-fusion bird populations [22] versus low inter-group movement rates by vervet monkeys [12] – and for various initial conditions.

In this study, we modelled a scenario in which two alternative behavioural preferences are introduced at the same time into a population of naïve individuals. This is consistent with cultural diffusion experiments. However, in natural settings, it is also likely that solutions to a foraging task might be discovered and rediscovered through repeated innovations [17]. Incorporating an asocial learning rate, whereby individuals can spontaneously learn to solve the puzzle using a certain solution, would be an interesting, and relatively straightforward, addition to our model. However, it should have very little impact unless it is large relative to the social learning rate, or if individuals do not abandon personal preferences to conform. Future research could also extend our model to reflect other characteristics. For example, including demographic processes could be a fruitful avenue for making long-term predictions. We assumed that the total carrying capacity of the environment had been reached and that each sub-population had a constant number of individuals. However, including varying population sizes could be interesting for exploring whether or not local traditions remain stable across multiple generations. Furthermore, the model may also be useful for considering how individual-level differences interact with the emergence and spread of culture. For example, juveniles could potentially learn faster than adults, or conformity could vary across age classes [18]. Individual-level differences have recently been highlighted as being important in shaping the dynamics of collective behaviour in animal groups [23,24]. It is therefore likely that such differences could play a major role in shaping the spread of behaviours and the establishment of local traditions in natural populations. It would also be interesting to consider a stochastic version of our model, since random events soon after traditions arrive in a naïve population are likely to play an important role in determining the tradition that ends up dominating.

In summary, our results provide new insights into the interplay between the movement of individuals and conformist learning in the emergence of animal culture. By simply incorporating these two processes, our model is able to make predictions about the emergence and stability of local traditions, and allow the influence of quantities such as initial population conditions and the degree of habitat fragmentation to be tested. A major strength of the model is its generalisability. Future research could extend the model to explore the spread of animal culture for more than two behavioural preferences, other environmental settings and different time scales, and integrating individual differences and non-static sub-population demographics. Such exploration of the spread of socially-transmitted information in animal populations has the potential to provide additional insights into the conditions under which local traditions emerge and persist.

## MATERIALS AND METHODS

### The baseline model

The spatially-explicit model describing the spread of animal culture integrates two processes: (1) transmission of information between individuals with a conformity bias, and (2) movement of individuals between spatially distinct patches of habitat. A novel behaviour, which consists of two equally difficult, and equally rewarding, solutions to a novel foraging resource (*s_1_* and *s_2_*) is introduced into a population of naïve individuals (who are unable to solve the task at the time of introduction) by adding innovators, which are individuals who know how to solve the task with a preference for either solution *s_1_* or *s_2_*. The information about how to solve the novel task, along with the preference for either one of the two alternative solutions, can be socially transmitted to other individuals. The spread of the two behavioural preferences in the population is then modelled, with simulations being run for 150 days (with a daily time step). At any time, an individual is either naïve, a solver *s_1_*, or a solver *s_2_*. During encounters with other individuals, naïve individuals can learn from solvers, and in doing so copy their behavioural preference, with parameter *α* governing the magnitude of the learning rate. The rate at which naïve individuals acquire one of the two alternative solutions (*s_1_* and *s_2_*) is a function of the proportion of solvers with this behavioural preference among all the solvers in the local sub-population. When individuals have a conformity bias (i.e. they are more likely to copy a specific behavioural preference than the prevalence of this preference among local solvers), which is given by the conformity parameter *λ*, then individuals use information about the behaviour of all other individuals in the patch when choosing which preference to acquire. The conformity parameter *λ* determines the strength of sigmoidality (i.e. S-shapedness) of the acquisition curve. An acquisition curve is the relationship between the prevalence of a preference for a solution in the local sub-population and the probability of adopting that preference (see equations below describing the conformist learning function *L* for learning *s_1_* and *s_2_*). In this model, conformist learning (from naïve to solver) and conformist switching (from solving the puzzle using one solution to using the alternative solution) were modelled in the same way using the same parameters. By doing this, the likelihood of an individual learning from another is approximately independent of whether or not the individual already has a preference for either solution to the puzzle.

In the initial baseline version of the model, the environment is assumed to consist of two patches, with each one hosting a sub-population. Within each sub-population, we assume individuals mix entirely at random. The size of each sub-population is at equilibrium throughout the simulation (i.e. no variation during the 150 days), essentially assuming that each sub-population size is at the strict carrying capacity of each patch. The movement rate of individuals between the patches decreases exponentially with the distance *d* separating them, with a parameter *m* determining the magnitude of the movement rate between the patches. In each patch *j* (where *j* = 1 or *j* = 2), the change of the numbers of individuals that are naïve (*U*^(*j*)^), solvers with a preference for solution 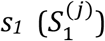 and solvers with a preference for solution 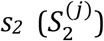 through time is modelled using a system of differential equations. For example, the change in the composition of individuals in patch *j* = 1 is given by:

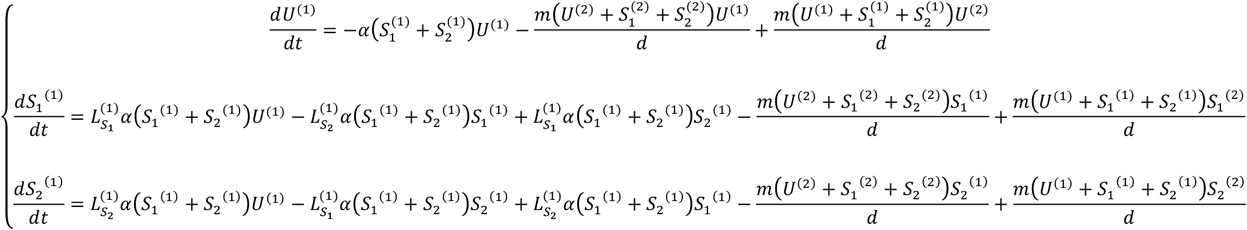

The equations for *j* = 2 are similar. In the equations above, the parameters 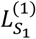 and 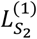 correspond to the conformist learning functions for learning solutions *s_1_* and *s_2_* respectively, which are functions of the prevalence of solution *s_1_* in the sub-population (*P*^(1)^; i.e. proportion of individuals in state *s_1_* among solvers in patch 1) and are defined as follows:

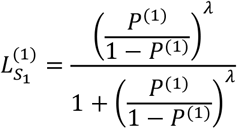

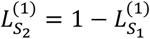

These conformist learning functions produce a sigmoidal relationship between a solution’s prevalence in the sub-population and the probability of adoption of that behavioural preference (called acquisition curve; [25,26]; Fig S4). If *λ* = 1, there is no conformity bias included in the model (i.e. straight 1:1 line; see Fig S4).

At the start of each simulation, two innovators (i.e. knowledgeable individuals) with solution *s_1_* were added to one patch and two innovators with solution *s_2_* were added to the other patch. We ran simulations for various conformity strengths: *λ* ∈ [1,5] sampled every 0.1; and movement rate magnitudes: *m* ∈ [0.0005,0.01] sampled every 0.0005. To investigate if results were affected by how quickly individuals learn, we ran the simulations for different learning rate: *α* = 0.001 (slow learning rate), *α* = 0.005 (intermediate learning rate) and *α* = 0.01 (fast learning rate). As there are only two patches here, changing the distance between the patches is equivalent to changing the movement rate *m* (see equations above), so we therefore set *d* = 1 for every simulation run. To investigate the effect of patch size, we also ran simulations with different numbers of naïve individuals in each patch at the start of the simulation (see Figs 1 and S1).

### Simple environmental setting

We extended the baseline model to three patches, each containing a sub-population. In this case, the equations described above for the baseline model were adapted for environmental settings with more than two patches (see Supporting Methods). At the start of each simulation, two innovators with solution *s_1_* were added to one patch and two innovators with solution *s_2_* were added to another patch (the third patch was assumed to initially consist of only naïve individuals). We ran simulations for the same ranges of values of conformity strength (*λ*) and movement rate (*m*) as for the baseline model. The learning rate was kept at the fast level (*α* = 0.01) for every simulation so that emerging patterns were more pronounced within the range of values explored for *λ* and *m*. We also investigated the effect of patch size by running simulations with the different numbers of naïve individuals in each patch at the start of the simulation (see Figs 2 and S2). To investigate the effect of habitat fragmentation, we varied the distance between the patch where two individuals trained to solve the puzzle with solution *s_2_* were added and the two other patches, investigating distances 1, 1.5 and 5, while the distance separating the two other patches was maintained at 1 (see schematics in Fig 2).

### Realistic environmental setting

Wytham Woods, Oxfordshire, UK (51° 46’ N, 01° 20’ W) is a 385ha broadleaf deciduous woodland surrounded by open farmland and covered by an evenly-spaced grid of 60 feeders (see the map in Fig S5). This is the location where Aplin et al. [11] performed the cultural diffusion experiment in great tits, introducing alternative novel foraging techniques and monitoring their spread. We extended the baseline model to this realistic setting, and used the adapted equations for more than two patches (see Supporting Methods). We started simulations with the same initial conditions as in the field study, releasing two innovators at targeted patches in a similar fashion (i.e. at the same feeders; see Fig S5). We divided the landscape so that each patch in our model contained one feeder. The total number of individuals across the woods and relative patch size (i.e. the number of individuals in each patch around each feeder in each time step) were derived from data described in [13]. We ran simulations for the same ranges of values for conformity strength as described for the baseline model. We investigated the following range of values for the movement rate: *m* ∈ [0.005,0.1]. This was different from the range of values explored for the two-patch and three-patch cases for this parameter because the distances separating patches were in meters here rather than in arbitrary spatial units. We modified the environment and the initial conditions to investigate how these changes affected the model outcomes. First, we used two distance measures between pairs of patches: *direct Euclidean distance* and *forest distance*, the latter being computed as the length of the shortest route between the two patches through the forest (without crossing open ground). This is known to be an ecologically relevant measure of distance with regard to movement within this population [13]. Second, we randomised the locations of feeders where trained individuals were released at the start of simulations. For three different values of conformity strength *λ* = 1 (no conformity included), *λ* = 1.2 (weak conformity) and *λ* = 4 (strong conformity), and a fixed movement rate magnitude (*m* = 0.02), we ran 100 simulations, each with a random location (i.e. sub-population/patch) where two innovators with solution *s_1_*, and another random location where two innovators with solution *s_2_*, were added at the start.

### Analysing emerging patterns

In all model runs for every environmental setting (two patches, three patches and Wytham Woods), we reported the total prevalence of solution *s_1_* across the whole population at the end of the simulation (*P_tot_*; i.e. proportion of individuals with behavioural preference for solution *s_1_* among all solvers in all patches) and the spatial variance of the final prevalence of solution *s_1_* in sub-populations (*P_var_*; i.e. variance in the proportion of individuals with behavioural preference for solution *s_1_* among local solvers in each patch). These two summary statistics were used to identify the emerging patterns:

- if *P_var_* > 0.1: *strong local traditions* established at the end of the simulation (i.e. some sub-populations are strongly dominated by one behavioural preference while the others are strongly dominated by the alternative preference)
- if 0.1 > *P_var_* > 0.01: *weak local traditions* established at the end of the simulation (i.e. some sub-populations have a bit more of one behavioural preference while the others have a bit more of the alternative preference)
- if *P_var_* < 0.01 and *P_tot_* > 0.66: *solution s_1_ dominated* across the whole system at the end of the simulation
- if *P_var_* < 0.01 and *P_tot_* < 0.33: *solution s_2_ dominated* across the whole system at the end of the simulation
- if *P_var_* < 0.01 and 0.33 > *P_tot_* < 0.66: *mixture of solutions* in every sub-population These criteria and thresholds were chosen in order to best reflect a visual identification of the emerging patterns (see examples in Fig 3).

